# Predicting changes in neutralizing antibody activity for SARS-CoV-2 XBB.1.5 using *in silico* protein modeling

**DOI:** 10.1101/2023.02.10.528025

**Authors:** Colby T. Ford, Shirish Yasa, Denis Jacob Machado, Richard Allen White, Daniel Janies

## Abstract

The SARS-CoV-2 variant XBB.1.5 is of concern as it has high transmissibility. XBB.1.5 currently accounts for upwards of 30% of new infections in the United States. One year after our group published the predicted structure of the Omicron (B.1.1.529) variant’s receptor binding domain (RBD) and antibody binding affinity, we return to investigate the new mutations seen in XBB.1.5 which is a descendant of Omicron. Using in silico *ico* modeling approaches against newer neutralizing antibodies that are shown effective against B.1.1.529, we predict the immune consequences of XBB.1.5’s mutations and show that there is no statistically significant difference in overall antibody evasion when comparing to the B.1.1.529 and other related variants (e.g., BJ.1 and BM.1.1.1). However, noticeable changes in antibody binding affinity were seen due to specific amino acid changes of interest in the newer variants.

## Introduction

In late November 2022, the United States Centers for Diseases Control stated that they began tracking a new SARS-CoV-2 variant known as XBB.1.5. At that time, XBB.1.5 was responsible for around 3% of all infections. Since then, XBB.1.5 has grown to represent 30% of all infections by January 2023 (1, 2).XBB.1.5 is characterized by 40 mutations in the Spike protein, 22 of which are in the receptor binding domain (RBD) (3). The highly prevalent mutations in the RBD are shown in Table 1 below.

**Table 1.** Prevalence of mutations in receptor binding domain of XBB.1.5. Prevalence is calculated as the percentage of samples (out of 10,168 from GISAID captured on February 12, 2023) that contain that mutation. Positions are numbered as their location in the larger Spike protein. These mutations are in comparison to the NC_045512.2 reference genome.

| Mutation | Sequences | Prevalence |
| --- | --- | --- |
| G339H | 10,000 | 98.35% |
| R346T | 9,992 | 98.27% |
| L368I | 9,839 | 96.76% |
| S371F | 9,857 | 96.94% |
| S373P | 9,856 | 96.93% |
| S375F | 9,860 | 96.97% |
| T376A | 9,852 | 96.89% |
| D405N | 9,895 | 97.32% |
| R408S | 9,616 | 94.57% |
| K417N | 9,329 | 91.75% |
| N440K | 9,687 | 95.27% |
| V445P | 9,653 | 94.94% |
| G446S | 9,673 | 95.13% |
| N460K | 9,757 | 95.96% |
| S477N | 9,984 | 98.19% |
| T478K | 9,960 | 97.95% |
| E484A | 9,943 | 97.79% |
| F486P | 9,919 | 97.55% |
| F490S | 9,927 | 97.63% |
| Q498R | 9,970 | 98.05% |
| N501Y | 9,983 | 98.18% |
| Y505H | 9,969 | 98.04% |

A health concern is that XBB.1.5 may evade existing antibodies derived from therapeutics, vaccination, and or previous Omicron (B.1.1.529) infection. It has been proposed that XBB.1.5 is a recombinant strain of the virus from BJ.1 and BM.1.1.1 as portions of the mutated Spike protein appear to be from each parent strain (4, 5). However, alternative hypotheses such as convergent evolution may also explain the similarity of portions of XBB.1.5’s mutated regions to those seen in other variants (6, 7).

In our previous work on the prediction of the receptor binding domain (RBD) structure of the Omicron variant, our process provided robust predictions, having a root mean square de-viation of atomic positions (RMSD) of 0.574Å between the predicted and empirically derived Omicron RBD structure (PDB: 7t9j) (8). Furthermore, our previous study proved useful as a predictive gauge of antibody efficacy several weeks prior to when empirical validations of the Omicron-antibody binding changes could be performed (9).

In this study, we use the methodology in our previous work to investigate XBB.1.5 and related variants (B.1.1.529, BJ.1, and BM.1.1) and expand the antibodies of interest to include more recently developed anti-Omicron antibodies. At the molecular level, we further elucidate the antibody binding and interfacing residues between three commercially-available antibodies: bamlanivimab, bebtelovimab, and tixagevimab and the RBD structure of each SARS-CoV-2 variant. We consider *in vitro* studies with existing antibodies and older variants to predict performance on XBB.1.5.

## Results

Given the four SARS-CoV-2 variants (B.1.1.529, BJ.1, BM.1.1, and XBB.1.5) and 10 antibodies, 40 *in silico* docking experiments were performed. As shown in Figure 1, the mean performance of the included neutralizing antibodies is similar across the four variants, with XBB.1.5 binding results being congruent to that of B.1.1.529. Furthermore, across the 10 antibodies tested, the binding affinities seen in XBB.1.5 are not weakened compared to B.1.1.529, nor are the differences statistically significant.

**Fig. 1.**
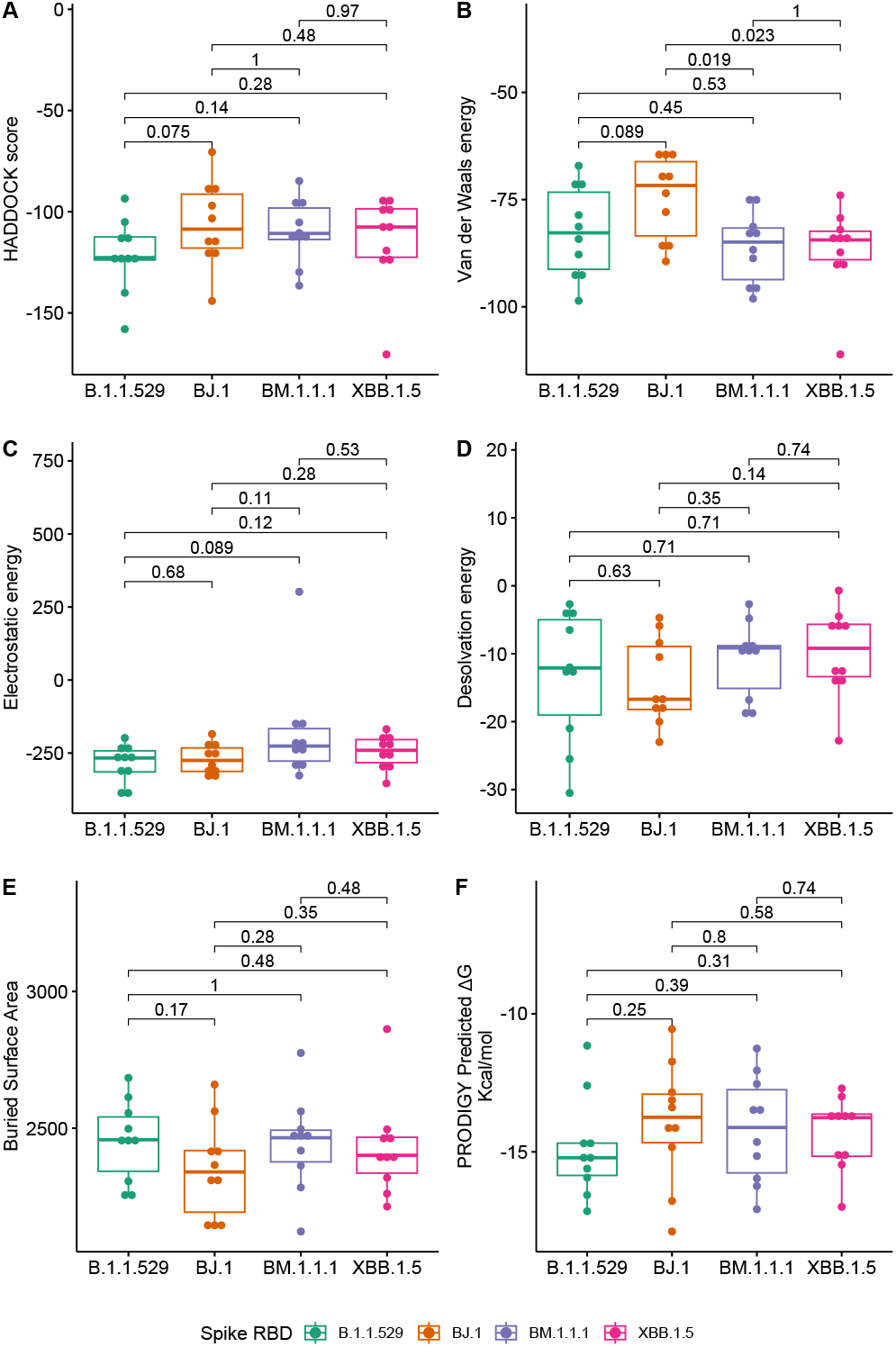
Boxplots of antibody binding performance by SARS-CoV-2 variant. Wilcoxon *p*-values are shown to assess the statistical significance of the differences between overall variant-antibody performance.

When assessing overall antibody performance against BJ.1, we see weakened Van der Waals energies as compared to the other three variants. This is depicted in the Uniform Manifold Approximation and Projection (UMAP) in Figure 2 where the position of some antibodies on the BJ.1 UMAP are increased (leading to decreased binding affinity). However, de-solvation energies and the buried surface areas are slightly improved overall when comparing BJ.1 results to the other three variant results.

**Fig. 2.**
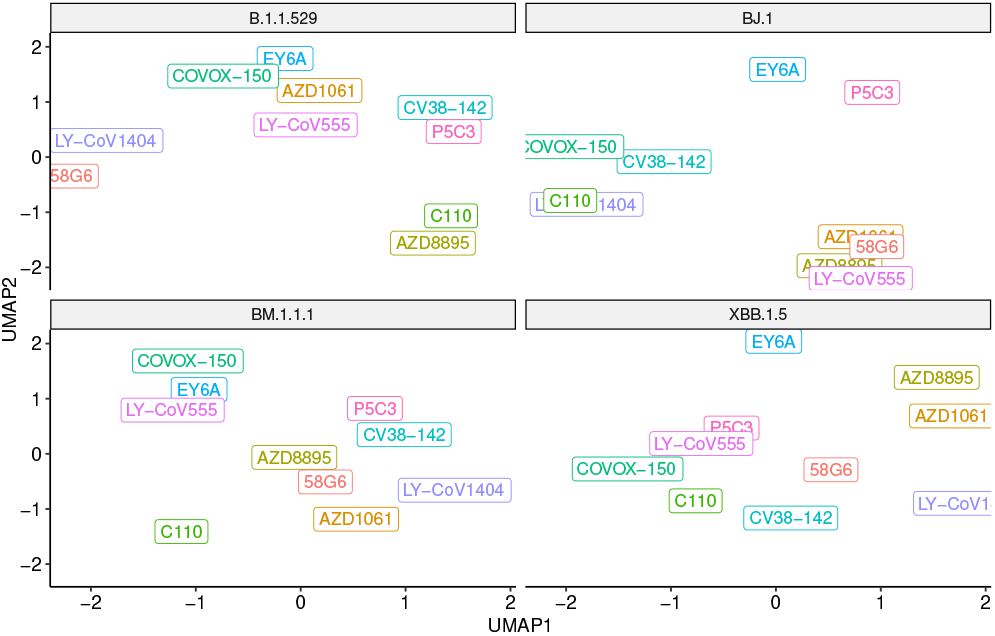
UMAP scatter plot of antibody binding affinity metrics with a variant in each quadrant. Note that a higher UMAP value is likely indicative of worse performance.

While there are instances of overall antibody performance increasing or decreasing in singular comparisons, we do not see an overarching pattern that indicates that XBB.1.5has evolved antibody evasion over B.1.1.529 (or BJ.1 and BM.1.1.1). In other words, XBB.1.5 does not appear to have evolved past current antibody defenses, specifically concerning the ten antibody structures tested in this study.

### Structural Changes in Antibody Binding Affinity

Of the ten antibodies tested in this study, we focus on the structural bases in which the antibodies LY-CoV555, LY-CoV1404, and AZD8895 work. The neutralization mechanisms of three antibodies have been extensively studied. These three antibodies have been available as therapeutics for treatment against COVID-19 infections (either currently or previously in the United States under Emergency Use Authorization) (10–12).

#### Bamlanivimab (LY-CoV555)

As shown in Figure 3, we see a consistent interaction between Bamlanivimab (LY-CoV555) and the variant RBDs at R/Q493. This differs from Jones et al. (13), which states that F490 and S494 in the RBD are the interfacing residues in this region.

**Fig. 3.**
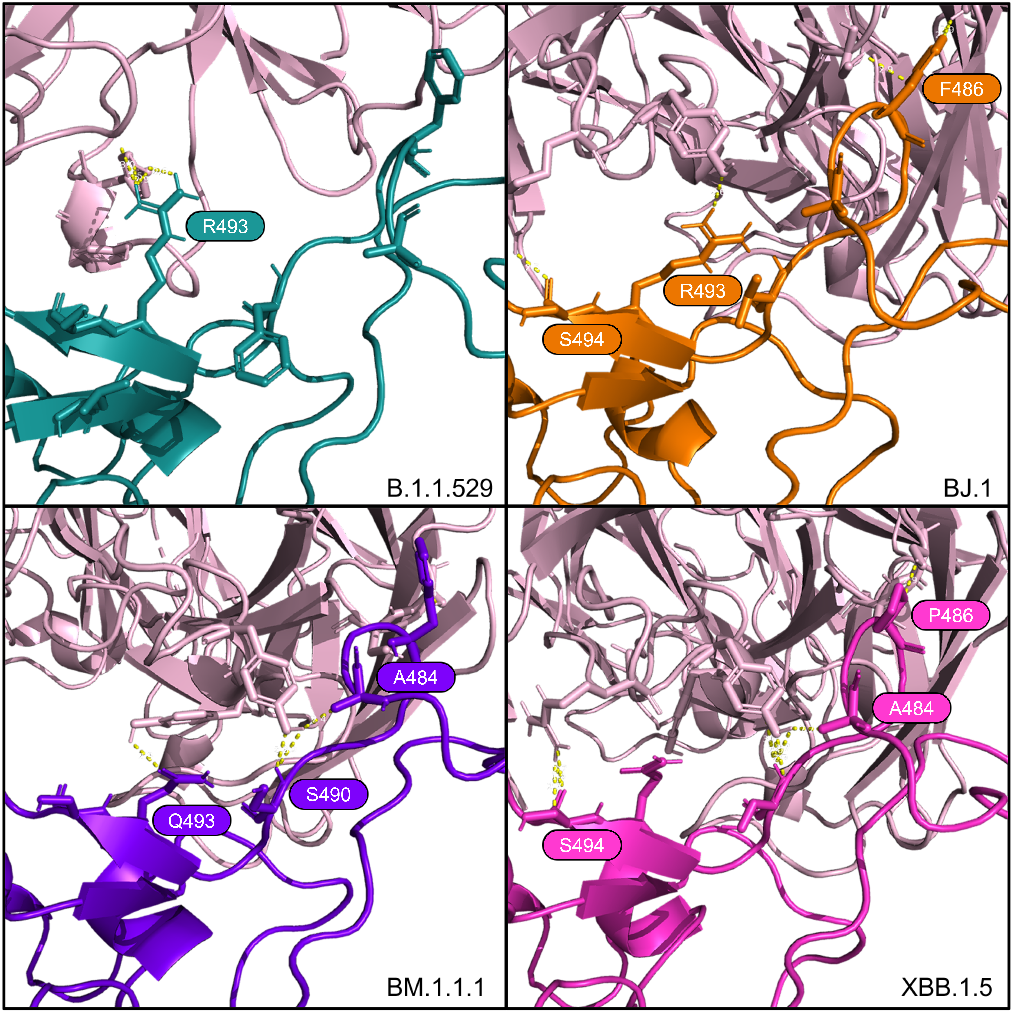
Interfacing residues of interest from the LY-CoV555 antibody (in pale pink) against the four RBD structures.

The PyMOL structural visualizations of the potential interaction residues coincides with the overall metrics returned from the HADDOCK analyses shown in Table 2. LY-CoV555 shows the worst overall performance against BJ.1 while the performance against B.1.1.529, BM.1.1.1, and XBB.1.5 are quite similar. Though not shown in Figure 3, the latter three complexes show a higher number of interfacing residues overall than in BJ.1, thus supporting the reported affinity metrics.

**Table 2.** Docking metrics for the LY-CoV555 antibody against the four RBD variants.

| Metric | B.1.1.529 | BJ.1 | BM.1.1.1 | XBB.1.5 |
| --- | --- | --- | --- | --- |
| HADDOCK score | -105.1±8.4 | -70.5±10.7 | -105.3±3.1 | -93.4±8.7 |
| Van der Waals energy | -87.8±5.1 | -65.1±6.2 | -95.3±6.1 | -87.3±2.6 |
| Electrostatic energy | -237.1±14.3 | -184.9±39.4 | -148.7±14.1 | -168.2±18.7 |
| Desolvation energy | -12±1.2 | -17.1±5.2 | -18.7±3.6 | -13±3 |
| Buried Surface Area | 2498.1±63.6 | 2155.5±97.3 | 2462.9±68.0 | 2386±52.3 |
| Average ΔG (σ) | -14.78 (0.13) | -14.16 (0.26) | -14.59 (1.15) | -14.21 (0.86) |

#### Tixagevimab (AZD8895)

For tixagevimab (AZD8895), as reported in Dong et al. (15), there is a critical contact residue at F486 on the RBD. We see this residue being interfaced in B.1.1.529, BJ.1, and BM.1.1.1. However, the F486 residue is mutated to proline at this position in XBB.1.5, though interactions from the antibody to the adjacent RBD residues at G485 and N487 of the RBD still occur. See Figure 5.

#### Bebtelovimab (LY-CoV1404)

Westendorf et al. (14) demonstrated that Bebtelovimab (LY-CoV1404) antibody binding affinity may not be affected by RBD mutations at E484, F490, Q493. Shown in Figure 4, we see consistent interactions from this antibody across all four variants around most of these positions in spite of mutations.

**Fig. 4.**
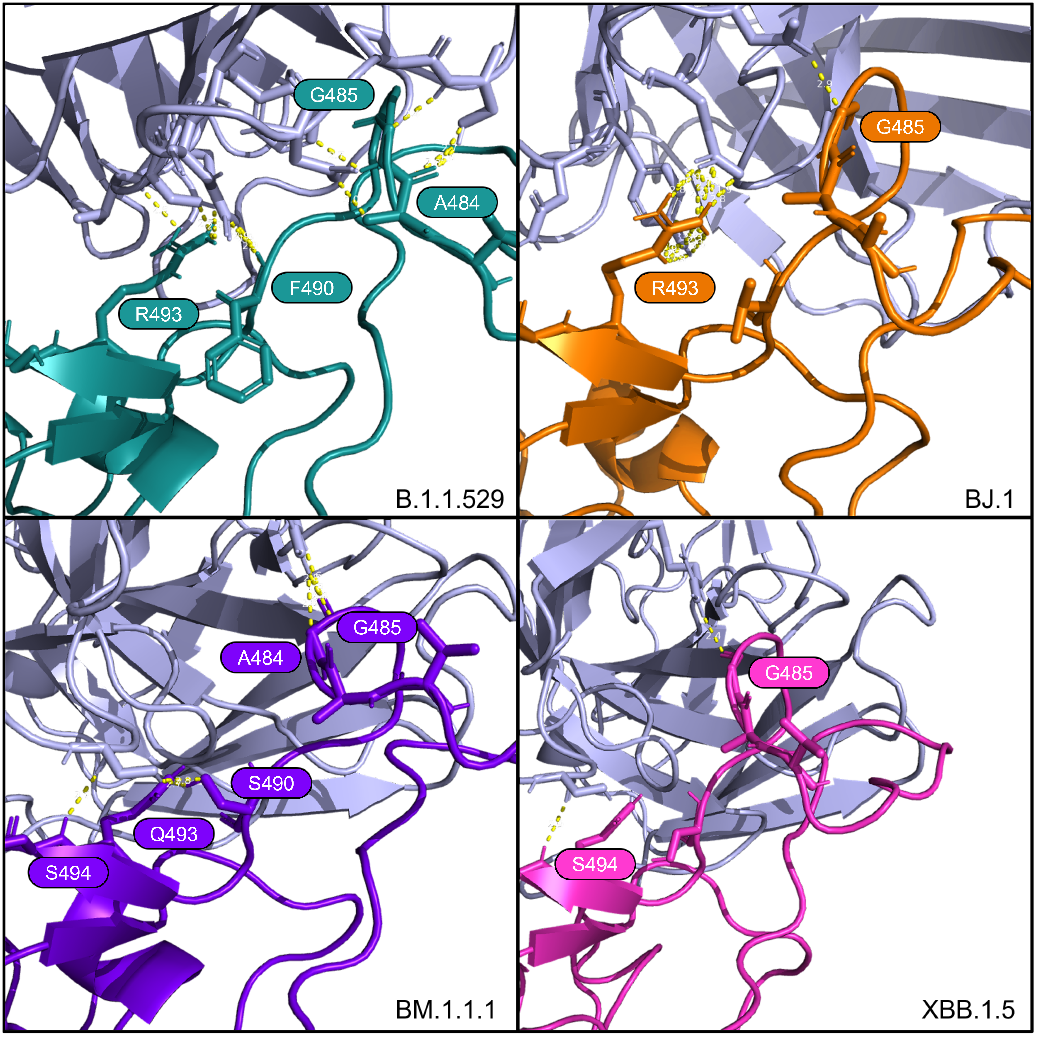
Interfacing residues of interest from the LY-CoV1404 antibody (in light purple) against the four RBD structures.

From the HADDOCK metrics shown in Table 4, this F486P mutation increases the binding affinity with the AZD8895, especially in terms of Van der Waals and electrostatic energies. Interfacing residues are abundant across all four of these AZD8895-RBD complexes (in addition to those shown in Figure 5), thus providing additional agreement to the strong affinity metrics reported by HADDOCK.

**Fig. 5.**
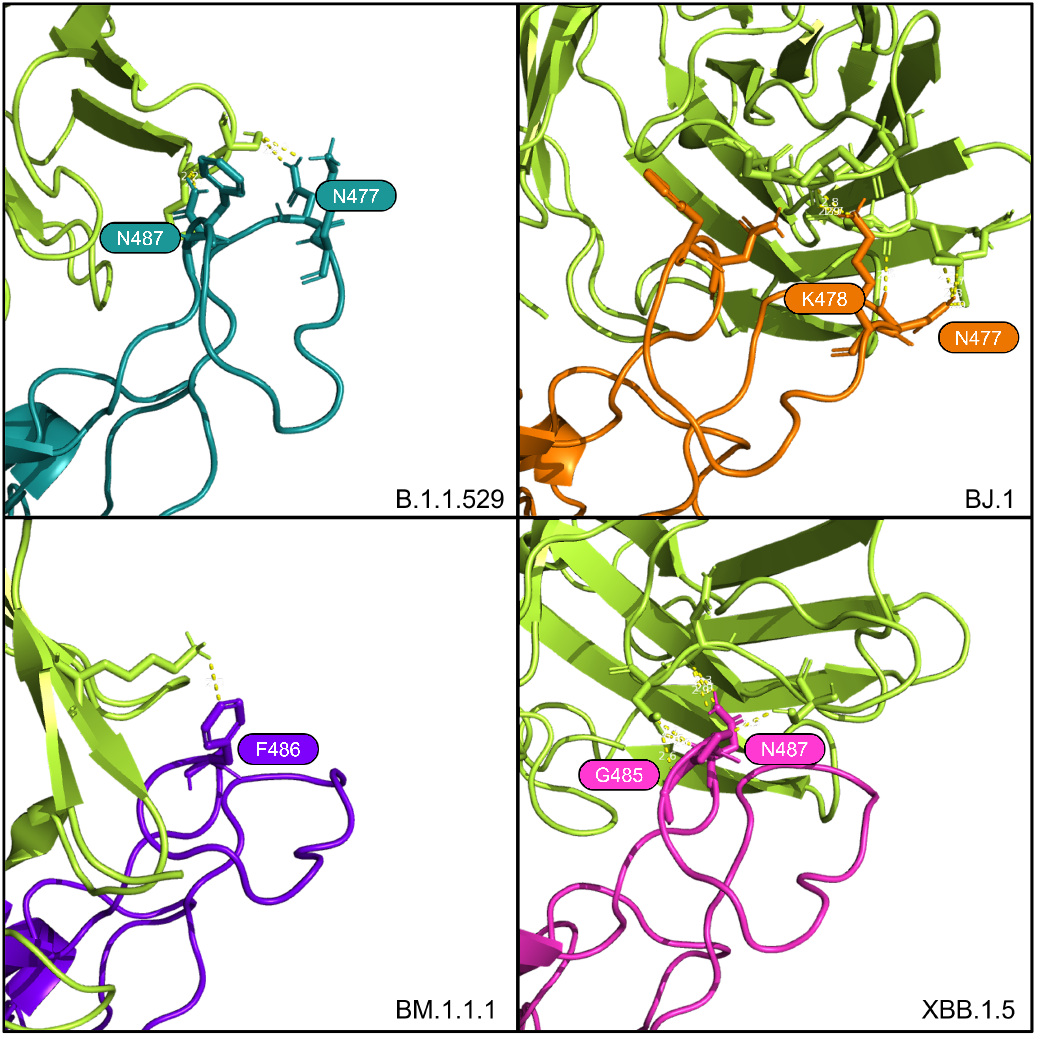
Interfacing residues of interest from the AZD8895 antibody (in bright green) against the four RBD structures.

These findings for LY-CoV1404 are congruent with the reported affinity metrics from the HADDOCK analyses shown in Table 3. Overall, HADDOCK scores are stable across the four variant complexes. The antibody LY-CoV1494 is predicted to have a slightly weaker interaction with XBB.1.5 compared to the other three variants.

**Table 3.** Docking metrics for the LY-CoV1404 antibody against the four RBD variants

| Metric | B.1.1.529 | BJ.1 | BM.1.1.1 | XBB.1.5 |
| --- | --- | --- | --- | --- |
| HADDOCK score | -124±1.1 | -115.4±1.8 | -110.9±6.3 | -109.1±8.5 |
| Van der Waals energy | -78.6±10.8 | -63.9±4.7 | -75.8±5.4 | -74±3.0 |
| Electrostatic energy | -262.1±24.3 | -337.7±27.6 | -290.7±16.1 | -291.8±7.3 |
| Desolvation energy | -30.5±7.1 | -23±6.2 | -2.7±2.8 | -4.5±3.5 |
| Buried Surface Area | 2457.2±111.6 | 2365.5±71.9 | 2467.1±80.9 | 2454.9±65.1 |
| Average ΔG (σ) | -16.47 (0.78) | -12.65 (0.53) | -19.90 (0.59) | -13.54 (0.88) |

## Methods

Our *in silico* modeling approach includes the curation or generation of the RBD structures for four SARS-CoV-2 variants and ten neutralizing antibody structures. Next, each antibody structure was docked against each RBD structure and binding affinity metrics were collected for comparison.

### RBD Structures

The Spike protein structure of the SARS-CoV-2 Omicron variant (B.1.1.529) was obtained from Protein Data Bank (PDB: 7t9j) (16). This structure was then trimmed to the RBD residues 339-528.

RBD sequences for BJ.1, BM.1.1.1, and XBB.1.5 were derived from representative samples found on GISAID:

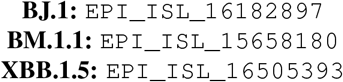

The RBD structures of BJ.1, BM.1.1.1, and XBB.1.5 were predicted with these sequences using AlphaFold2 (ColabFold-mmseqs2 version) (17, 18). Next, the most confident structure of each was used in docking analyses.

### Antibody Structures

Representative antibody structures were collected from various Protein Data Bank entries ranging from antibodies derived from infected patients (or patients with breakthrough infections) or commercially available antibodies used in the treatment of COVID-19. See Table 5.

**Table 4.** Docking metrics for the AZD8895 antibody against the four RBD variants.

| Metric | B.1.1.529 | BJ.1 | BM.1.1.1 | XBB.1.5 |
| --- | --- | --- | --- | --- |
| HADDOCK score | -93.5±4.3 | -88±6.7 | -84.8±3.2 | -123.8±6.9 |
| Van der Waals energy | -71.5±6.6 | -64.3±5.5 | -86.7±3 | -90.7±4.4 |
| Electrostatic energy | -230±29.9 | -215.5±5.9 | -150±11.3 | -256.3±222.6 |
| Desolvation energy | -4.5±1.5 | -8.4±5.7 | -8.9±1.6 | -5.5±2.8 |
| Buried Surface Area | 2259.5±49.1 | 2134.6±116.6 | 2482±46.4 | 2470.8±53.5 |
| Average ΔG (σ) | -12.55 (0.21) | -11.76 (1.78) | -13.08 (0.36) | -13.01 (0.77) |

**Table 5.** List of antibodies and their source PDB IDs used in this work.

| Antibody | Other Names | PDB ID | Citation |
| --- | --- | --- | --- |
| LY-CoV555 | bamlanivimab, LY3819253 | 7kmg | Jones et al. |
| LY-CoV1404 | bebtelovimab, LY3853113 | 7mmo | Westendorf et al. |
| AZD1061 | cilgavimab | 7l7e | Dong et al. |
| AZD8895 | tixagevimab | 7l7e | Dong et al. |
| 58G6 |  | 7e3l | Li et al. |
| CV38-142 |  | 7lm9 | Liu et al. |
| C110 |  | 7k8p | Barnes et al. |
| P5C3 |  | 7qtj | Fenwick et al. |
| EY6A |  | 7zf3 | Nutalai et al. |
| COVOX-150 |  | 7zf8 | Nutalai et al. |

Only a fragment antigen-binding (Fab) region of the antibodies was used in the docking analyses.

### Docking

To prepare the Fab structures, we renumbered residues according to HADDOCK’s requirements such that there are no overlapping residue IDs between the heavy and light chains of the Fab’s .PDB file. Residues in the Fab structures’ complementarity-determining regions (CDRs) were selected as “active residues” for the docking analyses.

Residues in the S1 position of the RBD were selected as the “active residues” of the RBD structures. Since all of the input RBD .PDB files were renumbered to numbers 339-528, all of the input RBD files share the same “active residue” numbers. Each of the ten antibody structures where docked against each of the four RBD structures using HADDOCK v2.4, a biomolecular modeling software that provides docking predictions for provided structures (27).

The HADDOCK system outputs multiple metrics for the predicted binding affinities and an output set of .PDB files containing the antibody docked against the RBD protein. PRODIGY, a tool to predict binding affinities using Gibbs energy, reported as ∆G in Kcal/mol units), was also run on each of the complexes (28).

This process resulted in forty sets of docked structures. Each set contains many antibody-RBD complex conformations, from which we selected the top-performing structure for each antibody-RBD pair. We used this top-performing complex for subsequent structural investigations into interfacing residues and docking positions.

These analyses were performed on the antibody-RBD structure pairs shown in Figure 1. The multiple metrics were used to assess the overall binding affinity changes between SARS-CoV-2 variants across multiple representative antibodies.

Further, the docked Protein Data bank Files (PDB) were manually reviewed using PyMOL (29) to search for interfacing residues and polar contacts between the RBD and Fab structures that may indicate neutralizing activity.

## Conclusions

Building on our previous work (8) in studying Omicron’s structure, we have continued to demonstrate the utility of *in silico* modeling for predicting whether antibody binding affinity changes with the evolution of new SARS-CoV-2 variants. Given that *in vitro* assessment of protein structure and antibody binding experiments are costly and take an extended time, *in silico* computational modeling provides a more economical and faster method near or at empirical resolution. Our previous *in silico* results were confirmed via an empirical study reported by VanBlargan et al. (9).

With computational modeling we rapidly preditct the potential severity of a new variant and provides predictions on antibody binding affinity. These predictions inform public health considerations and provide a method of rational drug design based on expected therapeutic and vaccine (and booster) efficacy. Computational modeling can be used to rapidly infer the public health consequences of a new variant in terms of the loss of efficacy of antibodies, such as breakthrough infections and associated healthcare burden.

For XBB.1.5 specifically, there are residue mutations of interest that may affect antibody binding of the various S1 region-binding antibodies tested here. Comparing previous reports on older variants concerning the three main antibodies discussed here to our computational results shows strong agreement between previous empirical results and our new *in silico* predictions.

For Bamlanivimab (LY-CoV555), Jones et al. (13) reported that mutations at RBD positions V483, E484, F490, and S494 either decrease or eliminate binding and function. Our study does not refute this, however, our computational modeling indicates the S494 is interfaced in the BJ.1 and XBB.1.5 interactions with LY-CoV555. This result suggests that this residue does not decrease the binding affinity of the antibody. Also, R/Q493 forms a polar interaction with LY-CoV555 in B.1.1.529, BJ.1, and BM.1.1.1, but not XBB.1.5.

In Bebtelovimab (LY-CoV1404), Westendorf et al. (14) states that mutations at E484 may confer advantages in antibody evasion capabilities for the virus. This result is supported in our study as A484 in our four variants in this study, is only interfaced in our docked structures for B.1.1.529 and BM.1.1.1. We see G485 as an important interfacing residue, forming a polar contact with LY-CoV1404 with all four RBD variants.

Lastly, Dong et al. (15) reports that aromatic residues from AZD8895 CDR loops form a hydrophobic pocket with the RBD residues G485, F486, and N487. Note that in XBB.1.5 there is a F486P mutation and, interestingly, the adjacent residues (G485 and N487) are interfaced in our predicted complex. It is possible the proline at position 486 provides less steric hindrance than phenylalanine, thus allowing surrounding residue interaction. This result can be tested in future studies.

The increased binding affinity of XBB.1.5 for ACE2 may lead to increased transmissibility at the population level (30). The results here do not indicate that we can expect increased disease severity on an individual level for patients that avail themselves of therapeutics and vaccination.

The climb in cases of COVID-19 disease linked to XBB.1.5 indicates that XBB.1.5 could be a very serious subvariant of Omicron. While other studies are needed to assess transmissibility, virulence, pathogenicity, and other facets of viral severity and epidemiology, this study predicts that many current therapeutic and infection-derived antibodies provide antibody binding affinities similar to B.1.1.529 for XBB.1.5. Thus, the results indicate that the health care outcomes should be positive for the patients that avail themselves of vaccines and therapeutics.

## Limitations and Future Work

This work estimates potential changes in antibody neutralization effects or antibody neutralizing affinity using *in silico* protein modeling and computational docking analyses. Given the computational and predictive nature of this study, empirical investigations are necessary to validate these findings. However, these computational approaches provide an economical, scalabl, and rapid methodology to understand the severity of new viral variants while the empirical work is being completed. Also, while HADDOCK is considered state-of-the-art in terms of protein docking, there are other docking tools that could pose different results for the comparisons

While we tested 10 representative neutralizing antibody structures against four variants of SARS-CoV-2, there are many more and antibody-variant RBD complexes to be assessed. In future work, we shall improve and automate our docking pipeline to enable to large-scale prediction of antibody binding affinity changes across any future SARS-CoV-2 variants of interest. Also, given the dynamic nature of protein structure conformations, alternative conformations may exist that show other polar contacts and antibody-RBD interfaces than those shown by the best performing HADDOCK complexes. All docking outputs and results, including those not shown in the body of this article, are available in the Supplementary Materials.

## Supplementary Materials

All code, data, results, docking parameters, and protein structure files can be found on GitHub at https://github.com/colbyford/SARS-CoV-2_XBB.1.5_Spike-RBD_Predictions.

## ACKNOWLEDGEMENTS

We acknowledge the following entities at the University of North Carolina at Charlotte: The Center for Computational Intelligence to Predict Health and Environmental Risks (CIPHER), North Carolina Research Center at Kannapolis, The Department of Bioinformatics and Genomics, The College of Computing and Informatics, and the University Research Computing group.

For the GISAID sequences EPI_ISL_16505393, EPI_ISL_16182897, and EPI_ISL_15658180, we thank the UNC Charlotte Environmental Monitoring Laboratory, the National Institute of Biomedical Genomics in India, and the British Columbia Centre for Disease Control Public Health Laboratory in Vancouver, Canada, respectively.

## Supplementary Note 1: RBD Structural Alignment and Comparison

Using PyMOL’s alignment tool (with 50 cycles and a cutoff of 2.0Å)(29), we superimposed the RBD structures as shown in Figure 6 and show RMSD metrics in Table 6.

**Fig. 6.**
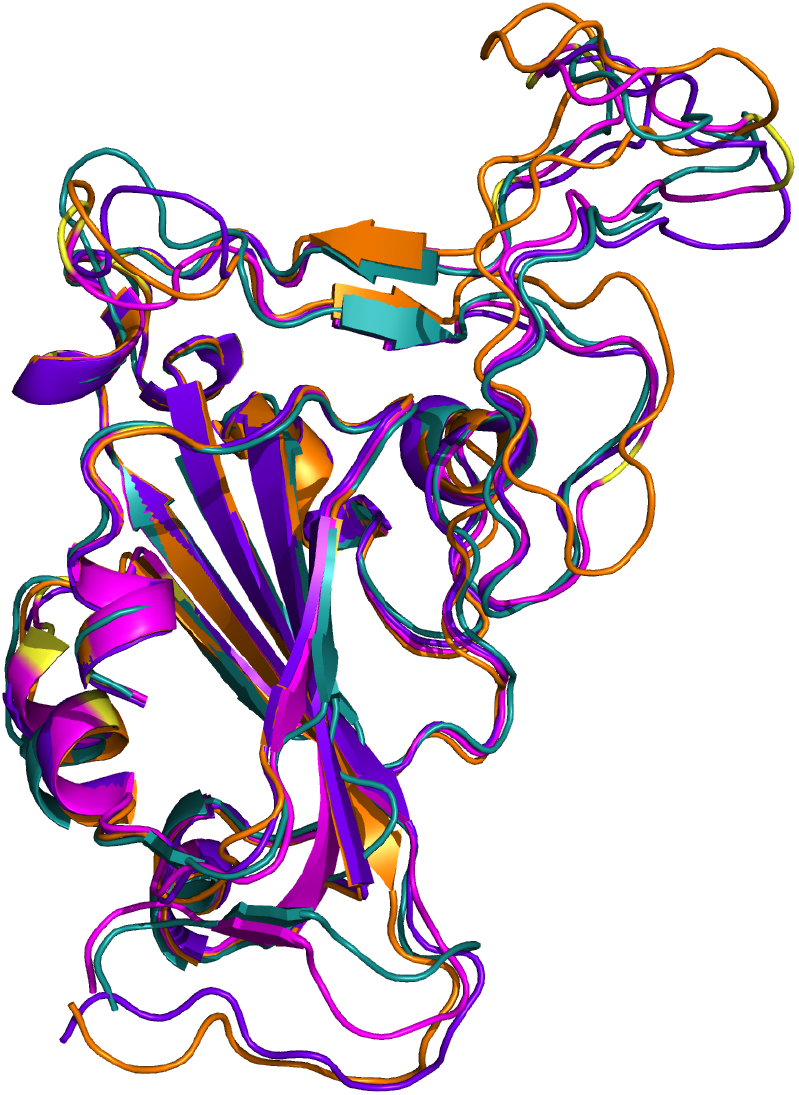
Cartoon representation of the four RBD structures aligned. B.1.1.529 (PDB: 7t9j) in teal, BJ.1 in orange, B.1.1.1 in purple, and XBB.1.5 in magenta with mutated residues from Table 1 highlighted in yellow.

**Table 6.** Alignment RMSD distance (in Ångstroms) between each variant’s RBD structure.

| Variant | B.1.1.529 | BJ.1 | BM.1.1.1 | XBB.1.5 |
| --- | --- | --- | --- | --- |
| B.1.1.529 | 0 | 0.780 | 0.708 | 0.653 |
| BJ.1 | 0.780 | 0 | 0.306 | 0.266 |
| BM.1.1.1 | 0.708 | 0.306 | 0 | 0.146 |
| XBB.1.5 | 0.653 | 0.266 | 0.146 | 0 |

## Supplementary Note 2: RBD Active Residue Predictions

Using CPORT, an interface predictor that provides a prediction of active and passive residues on a given protein (31), we evaluated the four RBD structures. This tool predicted that. See Table 7 below.

**Table 7.** CPORT-predicted active and passive residues of the four RBD structures.

| Variant | Active Residues | Passive Residues |
| --- | --- | --- |
| B.1.1.529 | 357, 359, 360, 361, 390, 391, 394, 417, 452, 453, 455, 456, 457, 470, 471, 476, 478, 479, 481, 482, 483, 484, 485, 486, 487, 489, 490, 492, 493 | 338, 351, 352, 355, 356, 362, 382, 383, 386, 388, 389, 396, 403, 409, 415, 416, 420, 421, 449, 450, 458, 459, 460, 461, 462, 465, 467, 468, 469, 472, 473, 474, 475, 477, 480, 494, 515, 516, 517, 518, 520, 521, 522, 523, 525, 526, 527, 528 |
| BJ.1 | 449, 450, 452, 455, 456, 457, 458, 459, 460, 461, 468, 469, 470, 471, 473, 474, 475, 476, 477, 478, 483, 484, 485, 486, 487, 489, 490, 493, 494 | 345, 346, 347, 351, 352, 403, 417, 419, 420, 421, 422, 444, 447, 448, 453, 462, 463, 465, 466, 467, 472, 479, 480, 481, 482, 495, 496, 497 |
| BM.1.1.1 | 338, 339, 357, 359, 360, 361, 362, 363, 364, 367, 388, 391, 392, 393, 394, 452, 455, 456, 472, 475, 483, 484, 485, 486, 487, 489, 490, 493, 494 | 340, 343, 344, 349, 351, 352, 355, 356, 366, 369, 370, 371, 372, 374, 382, 383, 384, 385, 386, 389, 390, 396, 403, 417, 421, 430, 448, 450, 457, 458, 459, 469, 470, 471, 473, 474, 476, 477, 478, 479, 480, 481, 482, 492, 495, 496, 497, 515, 516, 517, 518, 519, 523, 524, 525, 526, 527, 528 |
| XBB.1.5 | 351, 357, 358, 359, 360, 361, 390, 391, 393, 417, 421, 455, 456, 457, 483, 484, 485, 486, 487, 489, 493, 516, 518, 519, 520, 521, 522, 523, 525 | 338, 348, 352, 355, 356, 362, 364, 382, 383, 386, 388, 389, 396, 403, 409, 415, 416, 419, 420, 422, 430, 450, 452, 458, 459, 460, 461, 462, 465, 466, 467, 468, 469, 470, 471, 472, 473, 474, 475, 480, 490, 492, 514, 515, 517, 527, 528 |

Note that none of the mutations are predicted to disrupt the overall RBD tertiary structure in the AlphaFold2-generated structures. There are minor secondary structure changes as to where alpha helices or the anti-parallel beta sheet may begin or end, but overall the structures are very similar.

Looking at the main loop structure at the top of the S1 region, shown in Figure 7, the residue side chains are in similar positions, differing only by slight angular changes with the exception of the F486P mutation as previously mentioned. This proline mutation does not change the overall loop’s conformation, but provides rigidity at this location.

**Fig. 7.**
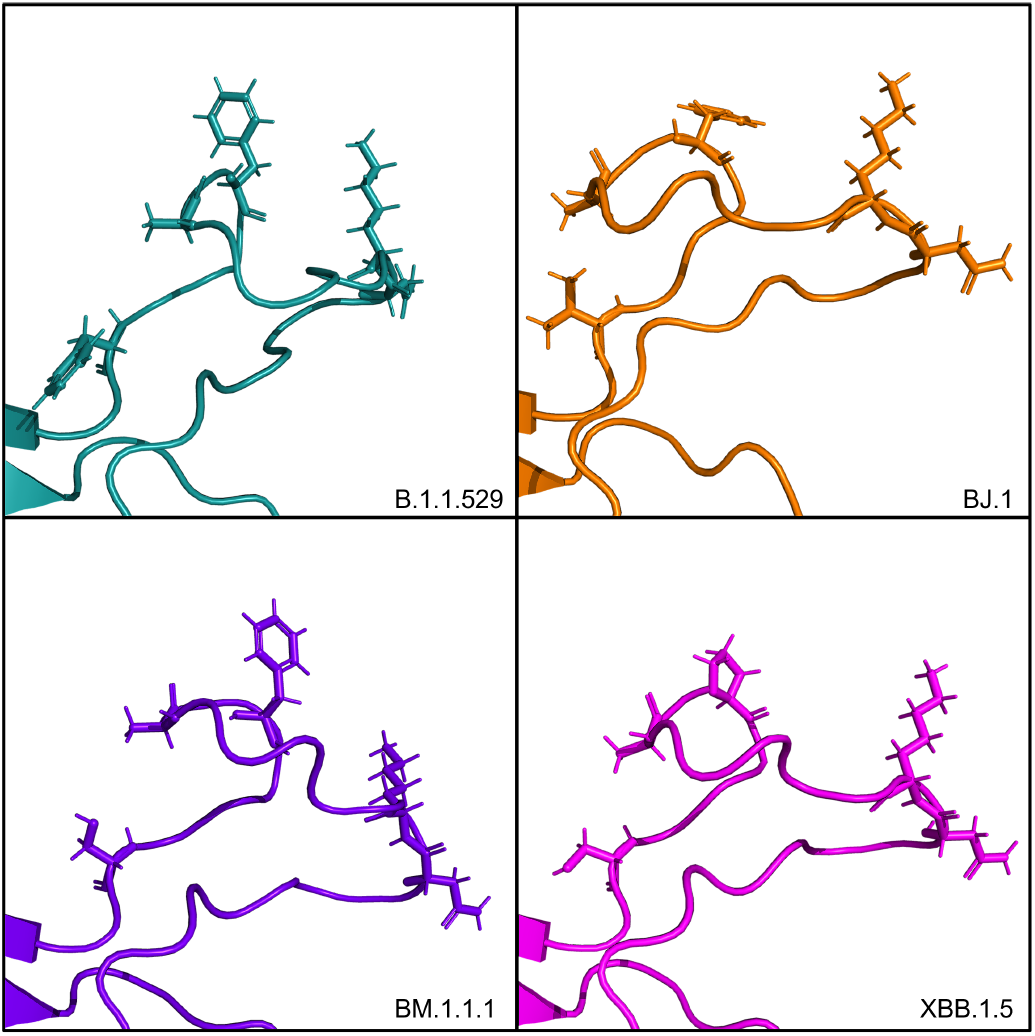
Side chains of the main loop at the S1 binding site for each of four RBD structures. Residues that are mutated from the wild type reference are shown as sticks.

Also, if we look at the residues that were predicted to be active visually, we can see that the majority of these residues concentrate around the S1 area of the RBD. Specifically, the loop structure on the top of the RBD that has been discussed heavily in this study and previous works is consistently predicted to contain multiple active residues across all four variant structures. See Figure 8.

**Fig. 8.**
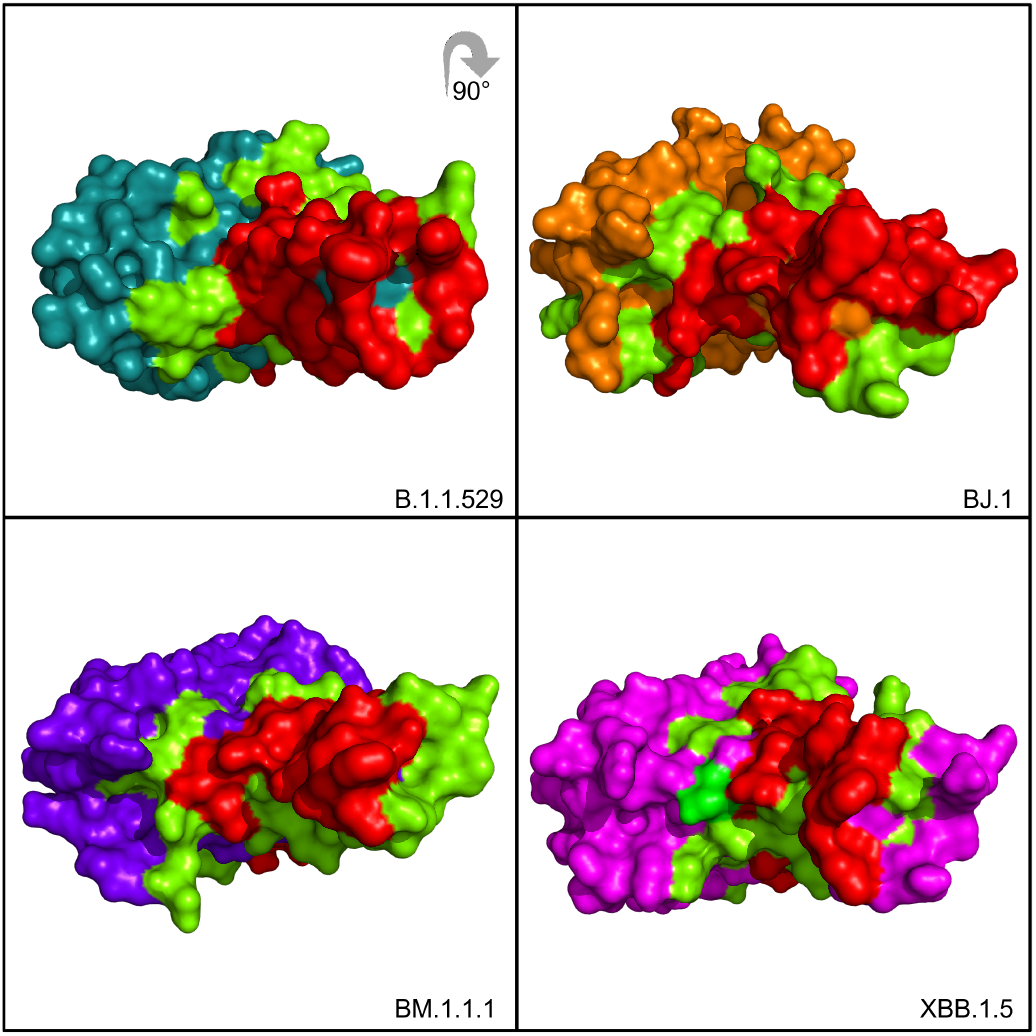
CPORT predicted active and passive residues (in red and green, respectively) of the four RBD structures. Shown as the “top” of the RBD, 90° foward.

## Supplementary Note 3: PRODIGY Results

In Figure 1, we show the top performing antibody-RBD complexes’ HADDOCK and PRODIGY scores. That is, the metrics for the top performing complex from the top performing HADDOCK cluster. In Figure 9 below, we show the individual PRODIGY scores of the top four complexes from the top performing HADDOCK cluster for each antibody-RBD experiment. There is agreement that no statistically significant drop in overall antibody binding affinity in terms of ∆G as predicted by PRODIGY.

**Fig. 9.**
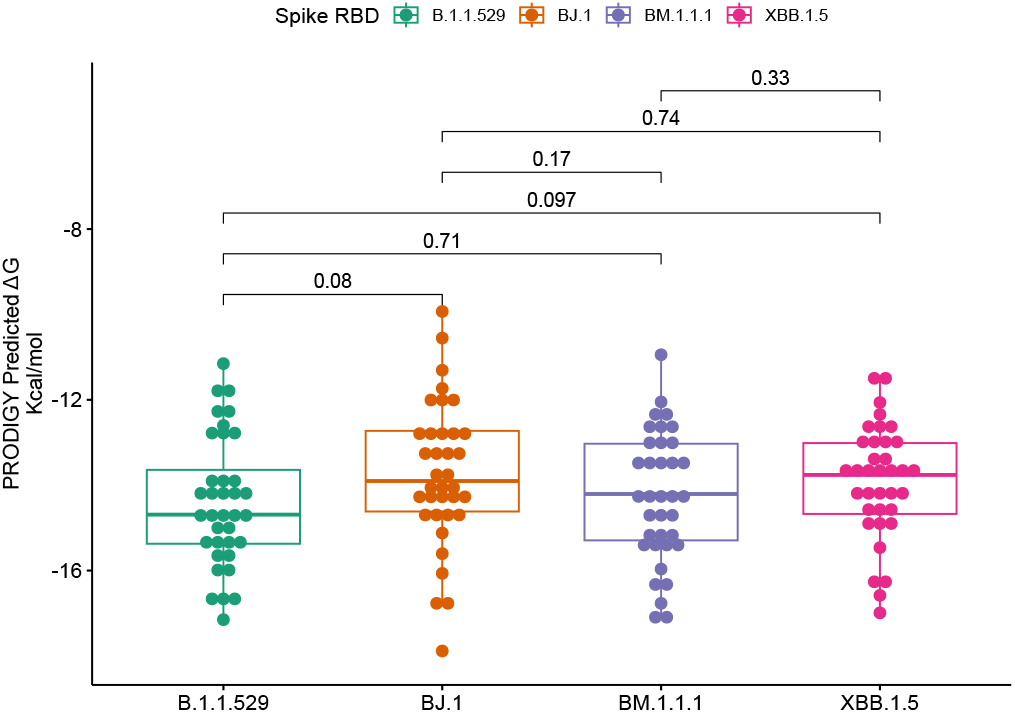
PRODIGY binding affinity metrics for the top four clusters in each antibody-RBD complex that resulted from the HADDOCK docking process.

There is considerable agreement between the CPORT predictions and the HADDOCK results listed in the main article. Nearly all of the interfacing residues detected in the complexes shown in Figures 3, 4, and 5 are predicted to be active residues from CPORT. Furthermore, many of these predicted active residues are also mentioned in Jones et al. (13), Westendorf et al. (14) and Dong et al. (15), thus further supporting that these residues on the top of the S1 region stand as the likely epitope between the RBD and various neutralizing antibodies evaluated in this study.

